# Naringenin as an antibacterial reagent controlling of biofilm formation and fatty acid metabolism in MRSA

**DOI:** 10.1101/2020.03.08.983049

**Authors:** Hun-Suk Song, Shashi Kant Bhatia, Ranjit Gurav, Tae-Rim Choi, Hyun Joong Kim, Ye-Lim Park, Yeong-Hoon Han, Jun Young Park, Sun Mi Lee, Sol Lee Park, Hye Soo Lee, Wooseong Kim, Yun-Gon Kim, Yung-Hun Yang

## Abstract

MRSA is Methicillin-resistant *Staphylococcus aureus* and they are widespread and making trouble in treatment in communities and surgical areas. MRSA have been adapted to antibiotics so that they can block the access of antibiotics physically or chemically deactivate it or modify the precursor of the target. Flavonoids are secondary metabolites which are naturally produced by plant or fungus and they are acting generally as pigment, quorum sensing molecules, antibiotics to other competitive microorganisms. Their natural origins and multiple activities have drawn much attention to be developed as the potential drugs since flavonoids could be a good candidate to overcome antibiotic resistant bacteria. Among various flavonoids, we found out naringenin has antibacterial activity on MRSA and Δ*agr* mutants which are more resistance than MRSA to beta-lactam antibiotics by decreasing biofilm formation dramatically and decreasing the secretion of fatty acid. It also showed high synergetic activity with oxacillin to both antibacterial activity and biofilm inhibition. Considering the number of flavonoids, our experiments expand the possibility of the use of flavonoids to MRSA.

## Introduction

MRSA stands for Methicillin-resistant *Staphylococcus aureus* and they are widespread and troublesome in the communities and hospital area (1). They occasionally cause pneumonia, urinary tract infection, surgical site infection, soft tissue infection and necrotizing pneumonia (2). The long-term overuse of antibiotics to treat them contributed to the multi-drug resistance in *Staphylococcus aureus* (3). MRSA have been developing their own way for antagonizing or blocking antibiotics. For instances, they have developed to have biofilm, peptidoglycan and membrane barrier to block the access of the antibiotics (4, 5). Additionally, antibiotics have been modifying targets of antibiotics to bypass them and chemical degradation or modifications of antibiotics have been made (6, 7).

Flavonoids are the secondary metabolites produced from plant and fungus and they fulfill various kinds of functions in them. For example, they act as a colorizing agent in plant and are involved in the filtration of UV (8). Additionally, they can be used as messengers as physiological regulators and cell cycle inhibitors in signal cascades in plant or fungus gene (9). Nowadays, flavonoids have drawn attention with their anti-bacterial activities to pathogenic bacteria (10). Since they derived from the natural origins, they seem safer to consumers than the chemically synthesized drugs. Also, they have advantages over antibiotics that they have multiple activities acting on the cells, so it is hard for bacteria to develop resistance to them (11). Thus, many approaches have been carried out to seek natural origin flavonoids having anti-bacterial effect and develop to make them have stronger efficacy (12). For example, catechins can disrupt the bacterial membrane by interacting with membrane and inhibiting enzymes (13). Isovitexin (apigenin-6-glycoside 14), epicatechin, and 5, 7, 5’-trihydroxyflavanol were active against *Staphylococcus aureus* having anti-biofilm activity (11). Naringenin, quercetin, sinensetin and apigenin have been shown as inhibitors of autoinducer-mediated cell-cell signaling (14). Synthetic 3-arylidene avanones acted as inhibitors of biofilm formation to *Staphylococcus aureus* and *Enterococcus faecalis* (15).

In this study, plant derived flavonoids including naringenin, chrysin, daidzein, genistein and apigenin were screened for the MRSA and MRSA mutant with higher resistance than USA300 (LAC). Naringenin decreased biofilm formation and fatty acid secretion was analyzed by GC-MS to see how they are distributed and affecting antibiotic resistance. We found out naringenin could decrease growth of MRSA, MRSA mutant and clinically isolated MRSAs decreasing biofilm formation at the same time. This showed the possibility of flavonoids as an antibacterial material or synergetic molecules with other well-known antibiotics.

## Result

### Flavonoids compound screening toward MRSA LAC and Δ*agr* mutant

Flavonoids have anti-bacterial activities to MRSA, and they can be simultaneously used with antibiotics for synergetic effect (16). And they have been tested for the MIC test toward pathogenic bacterium and suggested possible mechanisms but there are still detail explanations required to be discussed and the efficacy of the flavonoid are dependent on which bacteria they are applied to (17). Therefore, five flavonoids including naringenin, chrysin, daidzein, genistein and apigenin which have been previously studied to have been effective toward MRSA were selected to be tested for the Δ*agr* mutant with higher resistance. The chemical structures of five plant-derived flavonoids are presented in (Figure 1A). As an initial experiment, 100 μg/ml of flavonoids were applied to the LAC (WT) and Δ*agr* strain w/o or with oxacillin for synergetic effect confirmation by checking the comparative analysis of cell growth and biofilm formation.

**Figure 1.**
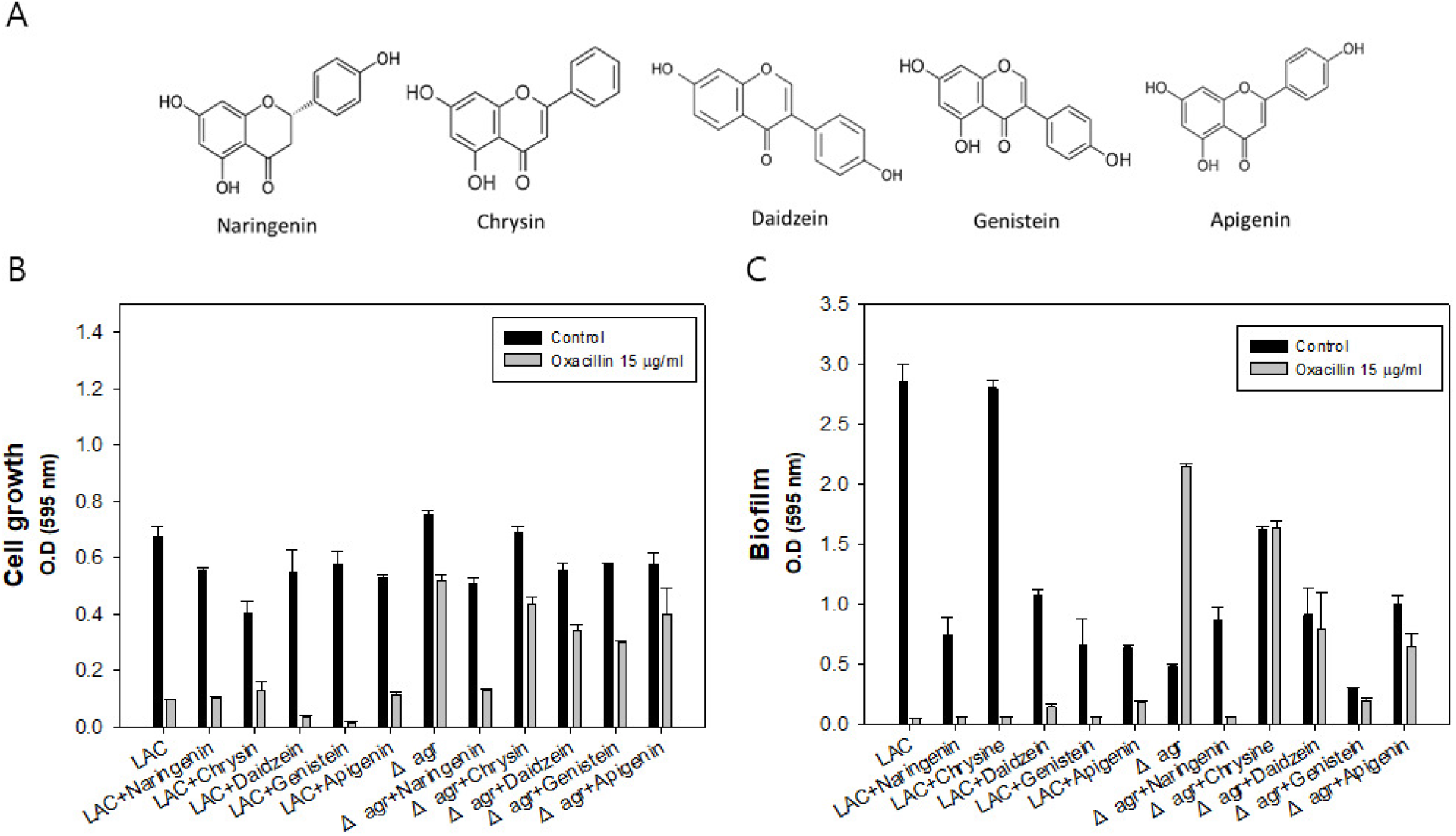
Chemical structures of plant-derived flavonoids, Investigation of five selected flavonoids efficacy to LAC and Δ*agr* strain. A: Cell growth of LAC and Δ*agr* strain with flavonoid w/o oxacillin or with oxacillin addition. B: Biofilm formation of LAC and Δagr strain with flavonoid w/o oxacillin or with oxacillin addition. The error bars represent the standard deviation of three replicates.

As a result, all the flavonoids were effective to LAC and Δ*agr* strain. Among them, the most potent efficacy was shown by naringenin with oxacillin in that the combination has synergistically decreased biofilm and completely inhibited the growth of both LAC and Δ*agr* strain (Figure 1B and 1C). Biofilm thickening is occasionally happening in the MRSA mutant with *agr*, and biofilm blocks the passage of the antibiotics thereby acquiring higher resistance. Thus, biofilm formation is one of the key factors affecting antibiotic resistance which should be dealt with when it comes to antibiotic treatment. Also, biofilm thickening was discovered especially when oxacillin was treated in Δ*agr* strain. In conclusion, the naringenin came out to be the most effective flavonoid to MRSA inhibiting growth and biofilm formation at this level though still mechanism and efficacy are dose dependent.

### Investigation of optimal concentration of naringenin and expression level of biofilm formatting genes

To set up the optimally effective concentration of naringenin to MRSA strains, different concentrations of naringenin were applied to MRSA strains. MIC_50_ value of the naringenin was similar between two groups so that 200 μg/ml of naringenin was used for further study unless stated otherwise. The problem of Δ*agr* strain is biofilm thickening when they grow since they lack PSMs which have surfactant activity causing lipid shedding and biofilm dispersion (5). Therefore, they have higher resistance to β-lactam anbitiotics. With addition of 200 μg/ml of naringenin with oxacillin, mutant strain was not able to grow, and they produce almost no biofilm (Figure 2).

**Figure 2.**
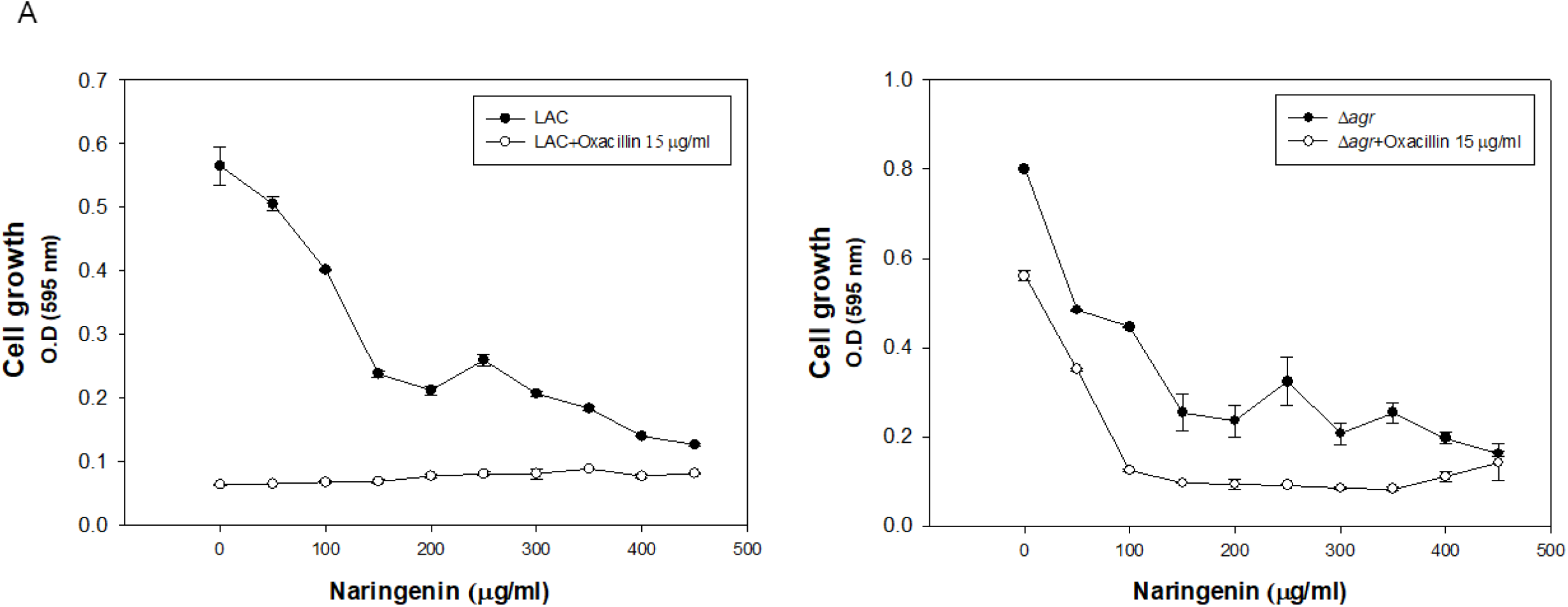
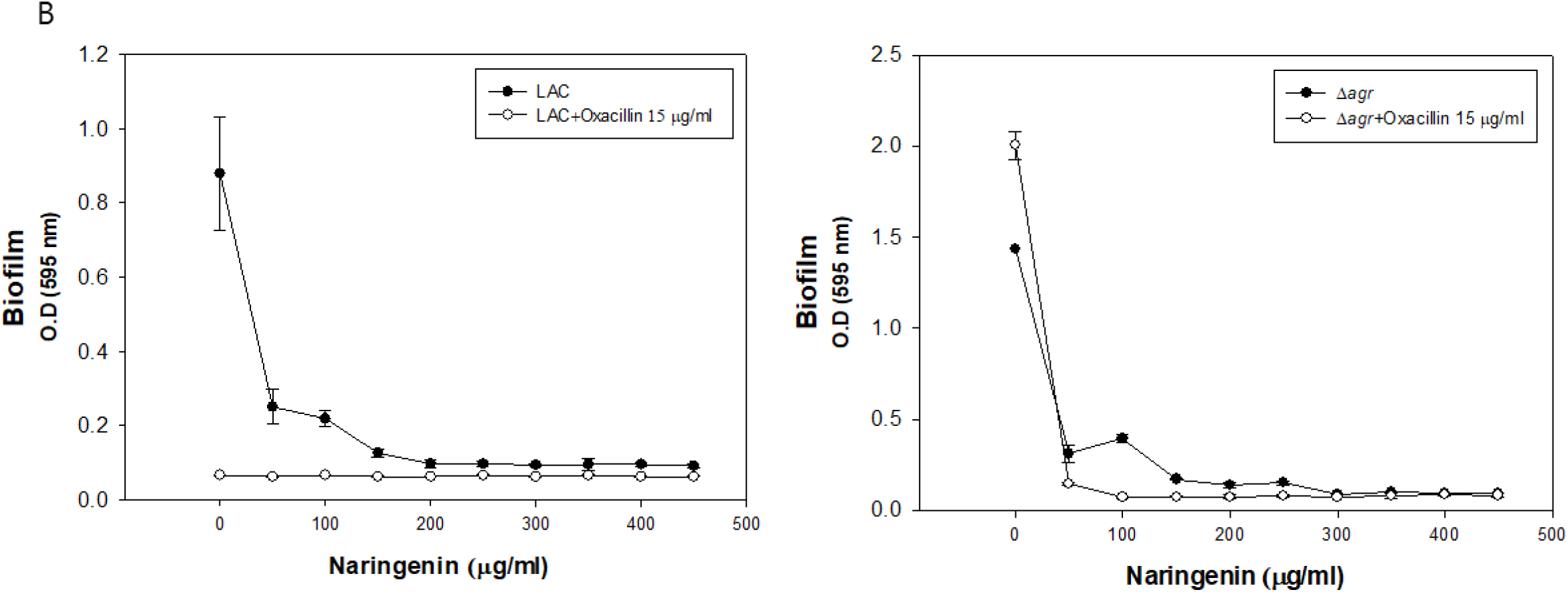
Investigation of optimal naringenin concentration to LAC and Δ*agr* strain. A: Cell growth of LAC and Δ*agr* strain with naringenin w/o oxacillin or with oxacillin addition. B: Biofilm formation of LAC and Δ*agr* strain with naringenin w/o oxacillin or with oxacillin addition. The error bars represent the standard deviation of three replicates.

To see any effect on mRNA expression, expression levels of *icaAD, mecA, nanK* were compared to see how they act on increased antibiotic susceptibility (Figure 3). *icaAD* genes are biofilm formation genes and *mecA* is a PBP2a protein coding gene increasing resistance of β-lactam anbitiotic. Finally, *nanK* is an indicator of persister cell formation so that the expression levels of between wild type and mutant groups were compared. In the case of *mecA* and *nanK*, there was no change in expression level with naringenin addition. When it comes to biofilm formation, the expression level of *icaAD* was less with naringenin addition compared to reference. As a result, with naringenin, decreased level of biofilm genes led to the higher susceptibility to oxacillin.

**Figure 3.**
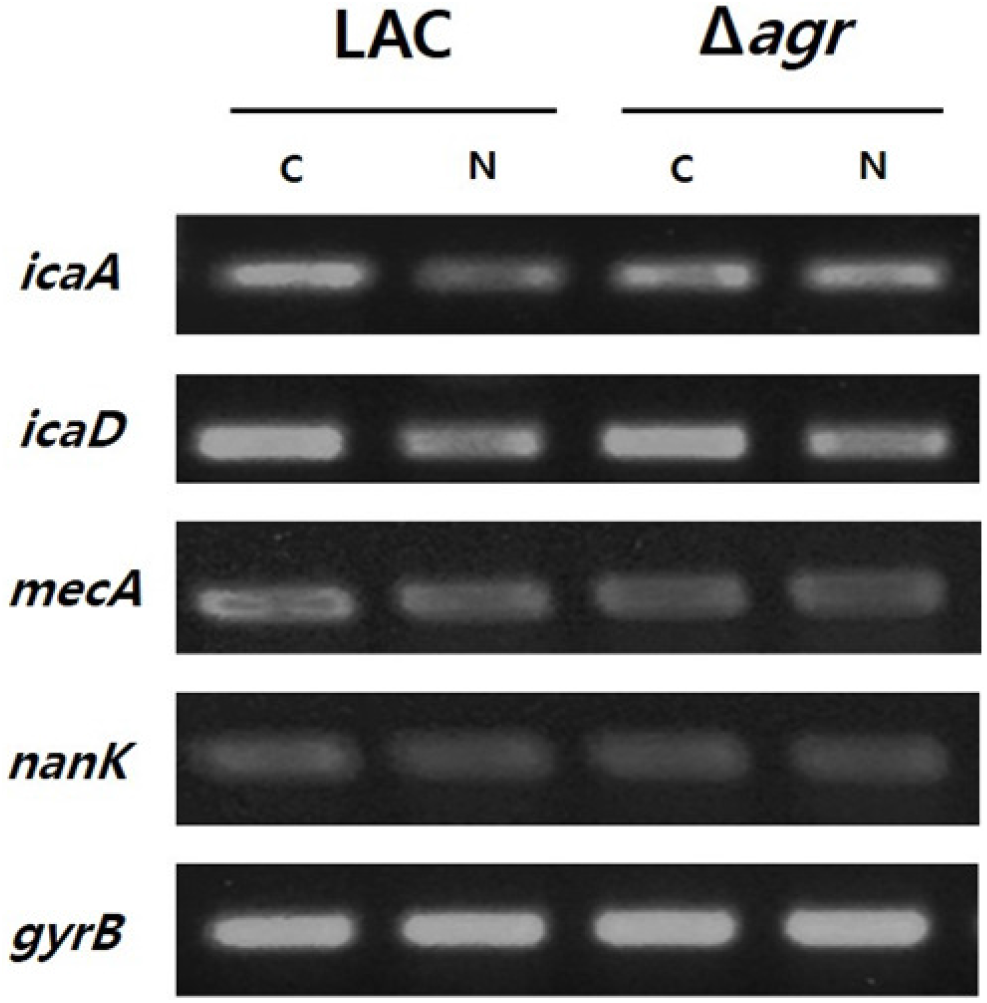
Semi-quantitative PCR of *icaAD, mecA* and *nanK*.

### Monitoring of synergetic effect by naringenin

To see the synergetic effect of naringenin, we applied oxacillin, naringenin and combination of two to both LAC and Δ*agr* mutant. In LAC strain, naringenin showed antibacterial activity resulting the decrease of growth and biofilm formation (Figure 4A). In Δ*agr* mutant, the effect of naringenin was critical this is because the expression of *icaAD* was higher in Δ*agr* mutant than LAC with naringenin treatment. Combination therapy showed higher antibacterial activity than oxacillin alone and its activity on biofilm inhibition was much higher than oxacillin alone (Figure 4B). Considering oxacillin increasing the production of biofilm, the effect of naringenin to decrease the biofilm was quite useful and its combinatorial use to Δ*agr* mutant clearly showed the dramatic decrease of biofilm formation with antibacterial activity. When we monitored biofilm formation with SEM at different time points by treating oxacillin, naringenin and both, we could see the clear differences. As detected by absorbance of growth and biofilm formation, SEM data showed the less formation of biofilm and low and dispersed population of Δ*agr* strains by treating both oxacillin and naringenin (Figure 5A, 5B). As a result, naringenin could work as a sole antibiotic material and anti-biofilm material, but it could be also used as a synergetic molecule with known antibiotic, oxacillin.

**Figure 4.**
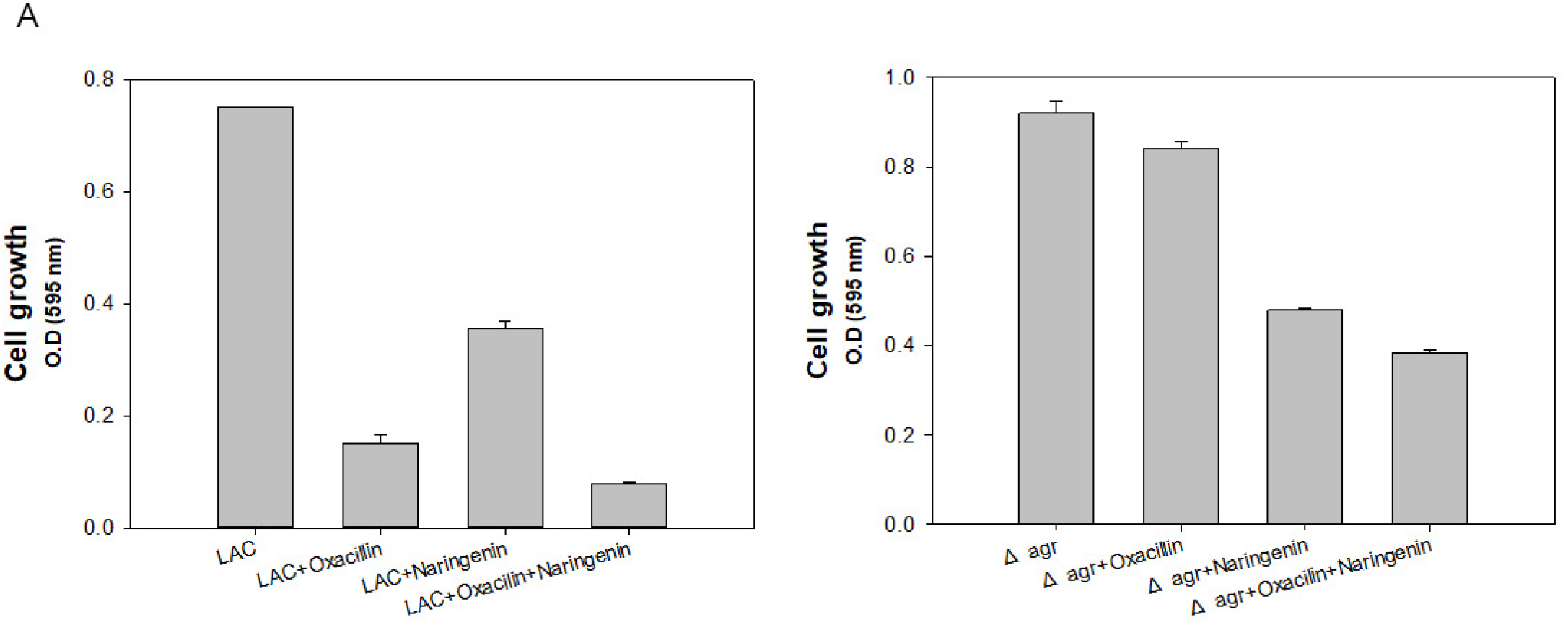
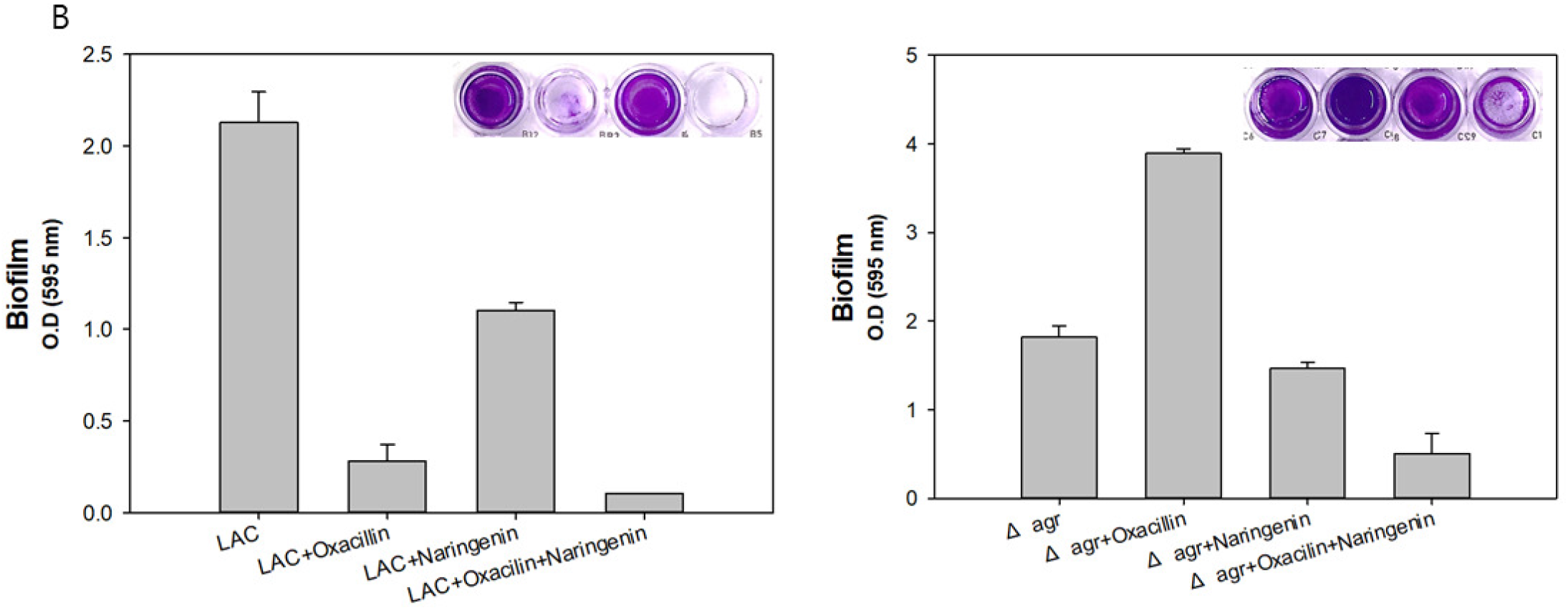
Growth and biofilm formation profile of MRSA with higher resistance compared to LAC. A: Cell growth of LAC and Δ*agr* strain with naringenin w/o oxacillin or with oxacillin addition. B: Biofilm formation of LAC and Δ*agr* strain with naringenin w/o oxacillin or with oxacillin addition. The error bars represent the standard deviation of three replicates.

**Figure 5.**
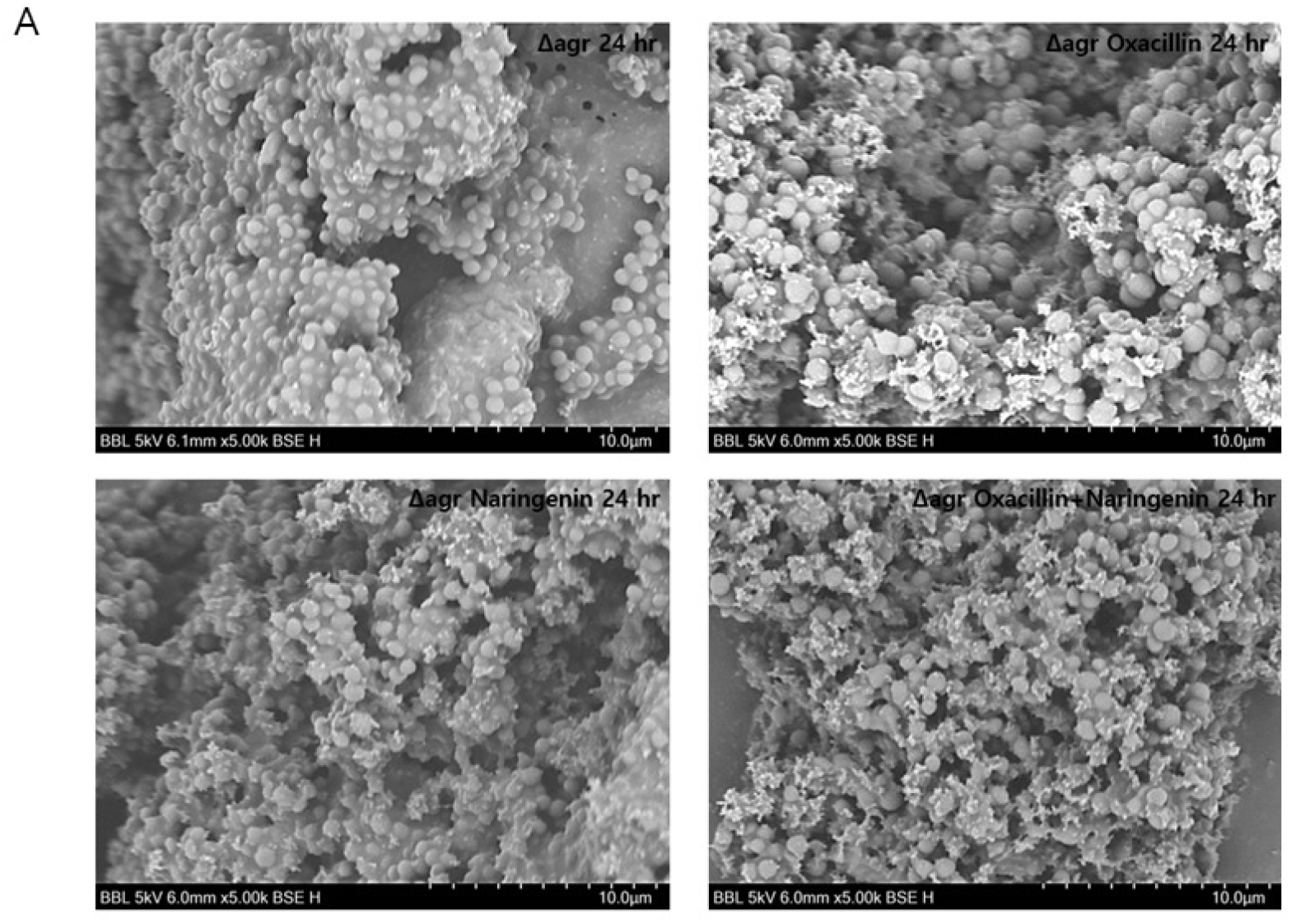
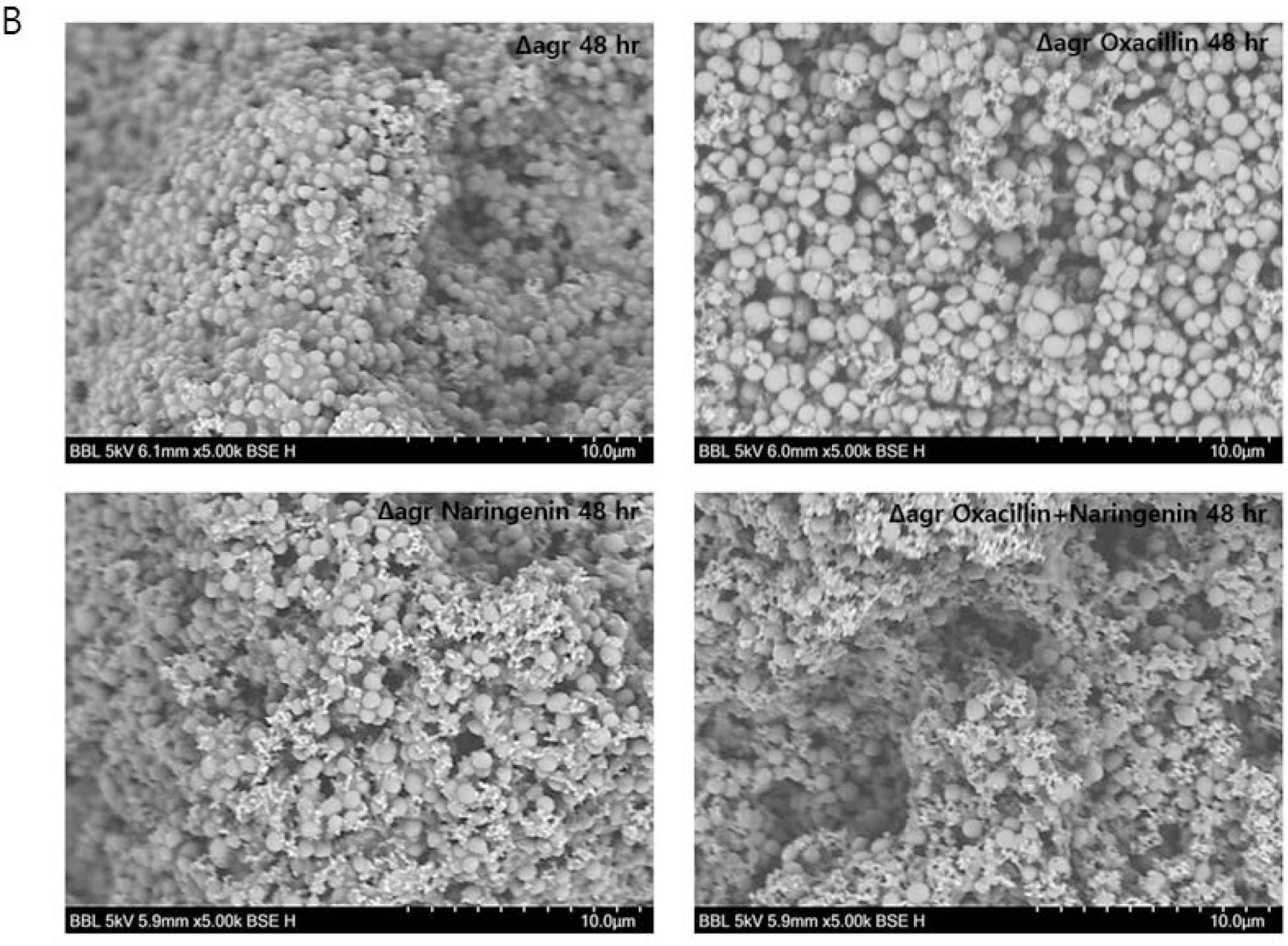
Scanning electron microscopy (SEM) images of with or w/o oxacillin and naringenin addition. A: SEM images of Δ*agr* strains with or w/o oxacillin and naringenin treatment at 24 hr. B: SEM images of Δ*agr* strains with or w/o oxacillin and naringenin treatment at 48 hr.

### Changes in fatty acid metabolism and expression level of biofilm formation genes

Naringenin has been known to be inhibit quorum sensing limiting biofilm formation and can reduce fluidity of the bacterial membrane (18). In addition to that, naringenin can target FAS II system which is essential pathway for bacteria survival.

In Δ*agr* mutant there was lower level of secreted fatty acid since fatty acid are known to have anti-bacterial and anti-biofilm activities and external addition of fatty acid increase the antibacterial activity and antibiofilm activity (19). Thus, it is favorable for them to keep lower amount of especially C16 and C18 fatty acids for their own survival. Microenvironment control of MRSA is quite related to control of antibiotic resistance so that fatty acid distribution profile of mutant strains was conducted w/o or with naringenin addition. We performed the analysis of secreted fatty acid at different time points with two LAC and Δ*agr* strains. Preparation of materials were explained in Material and methods section. Surprisingly, with naringenin addition, total amount of fatty acids was decreased in both strains (Figure 6A). Although total amount of fatty acids by cell weight was increased in LAC with naringenin, but there was growth limitation so that it is out of consideration (Figure 6B). Also, total fatty acids production was lower when treated naringenin (Figure S1). Considering flavonoids are the inhibitor of fatty acid synthesis, and secreted fatty acid could affect antibiotic activity, this is quite interesting result and explained the functional mode of naringenin. This have led to the higher effect of oxacillin and lower level of biofilm formation to kill both LAC and Δ*agr* strains.

**Figure 6.**
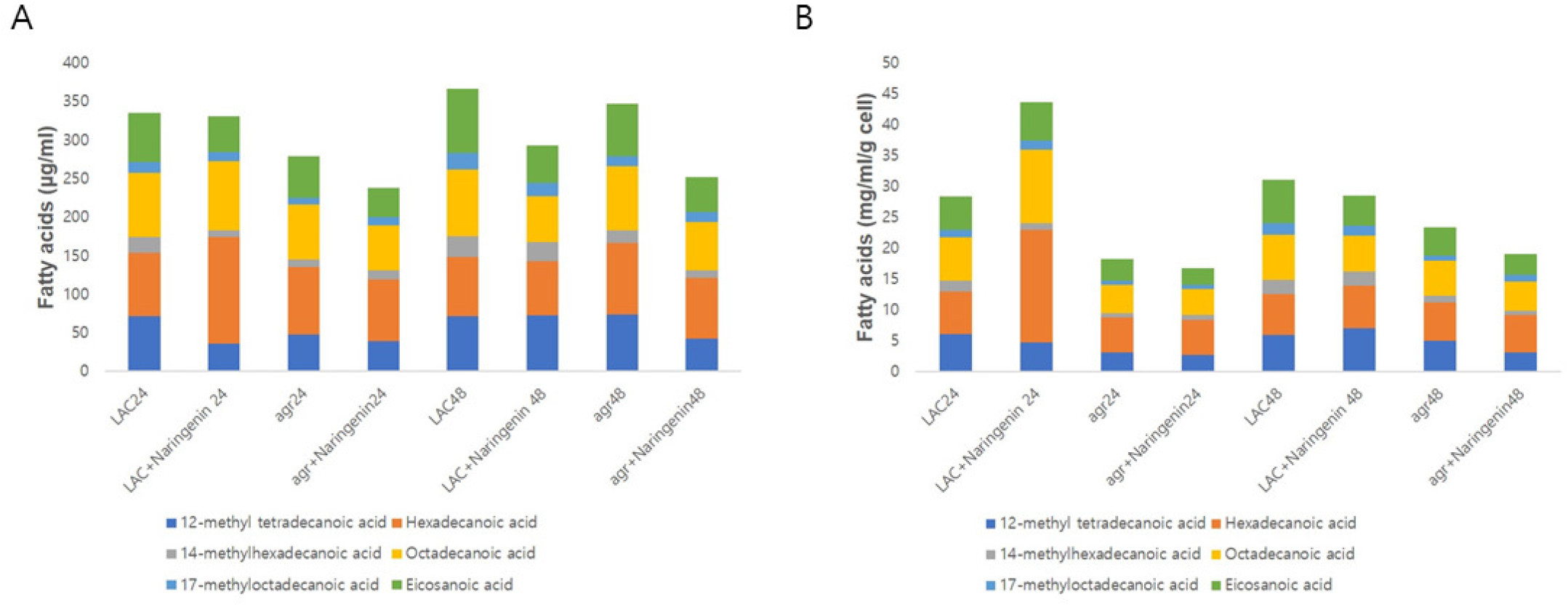
Analysis of secreted fatty acid w/o or with naringenin in MRSA mutants. The error bars represent the standard deviation of three replicates.

### Application of synergetic effect of naringenin and oxacillin to clinically isolated strains

Naringenin and oxacillin had synergetic effect toward LAC and Δ*agr* but still they can be classified as CA-MRSA which has MIC level about >32 μg/ml. Thus, we further explored the effect of combination of naringenin and oxacillin to 10 clinically isolated MRSAs. The concentrations of naringenin and oxacillin were set same for the comparison to CA-MRSA data. Surprisingly, the synergetic effect of the oxacillin and naringenin still remained in the most of clinically isolated strains except MRSA14459 (Figure 7A). Oxacillin alone was able to treat MRSA14459 but not with naringenin combination. In the case of biofilm, all the strains did not produce biofilm when treated with oxacillin and naringenin simultaneously (Figure 7B). To validate the case of MRSA14459, we further investigated the MIC of each strains. However, in this time, MRSA14459 was able to tolerate the antibacterial effect of oxacillin even over 200 μg/ml. Except MRSA 6230 of which MIC is about 150 μg/ml, MIC level of oxacillin of other strains were over 200 μg/ml (Table 1). Inequal effect of oxacillin might be from induction of persister formation or from unknown mechanism (data not shown). In addition, it was surprising that even MRSA6230 and MRSA14459 have higher resistance to oxacillin since they are SCCmec type IV which is a characteristic of CA-MRSA. However, synergetic effect of oxacillin and naringenin still exist even at the low concentrations of oxacillin for both strains. Therefore, utilization of naringenin and oxacillin together is an effective way to treat most of the MRSAs though the range of concentration should be increased depending on strains for optimal combination therapy.

**Tab_1.** Types of clinically isolated MRSAs and Oxacillin MIC for each strains.

| Name | Type | SCCmec | Oxacillin<br>MIC in µg/ml | spa type | MLST ST |
| --- | --- | --- | --- | --- | --- |
| LAC | MRSA | IV | 20 | t008 | 8 |
| 2065 | MRSA | III | >200 | t037 | 239 |
| 6230 | MRSA | IV | 150 | t324 | 72 |
| 6288 | MRSA | III | >200 | t037 | 239 |
| 7557 | MRSA | II | >200 | t9353 | 5 |
| 7875 | MRSA | IV | >200 | t664 | 72 |
| 8471 | MRSA | II | >200 | t9353 | 5 |
| 9291 | MRSA | II | >200 | t601 | 5 |
| 12779 | MRSA | II | >200 | t2460 | 5 |
| 14278 | MRSA | II | >200 | t9353 | 5 |
| 14459 | MRSA | IV | >200 | t324 | 72 |

**Figure 7.**
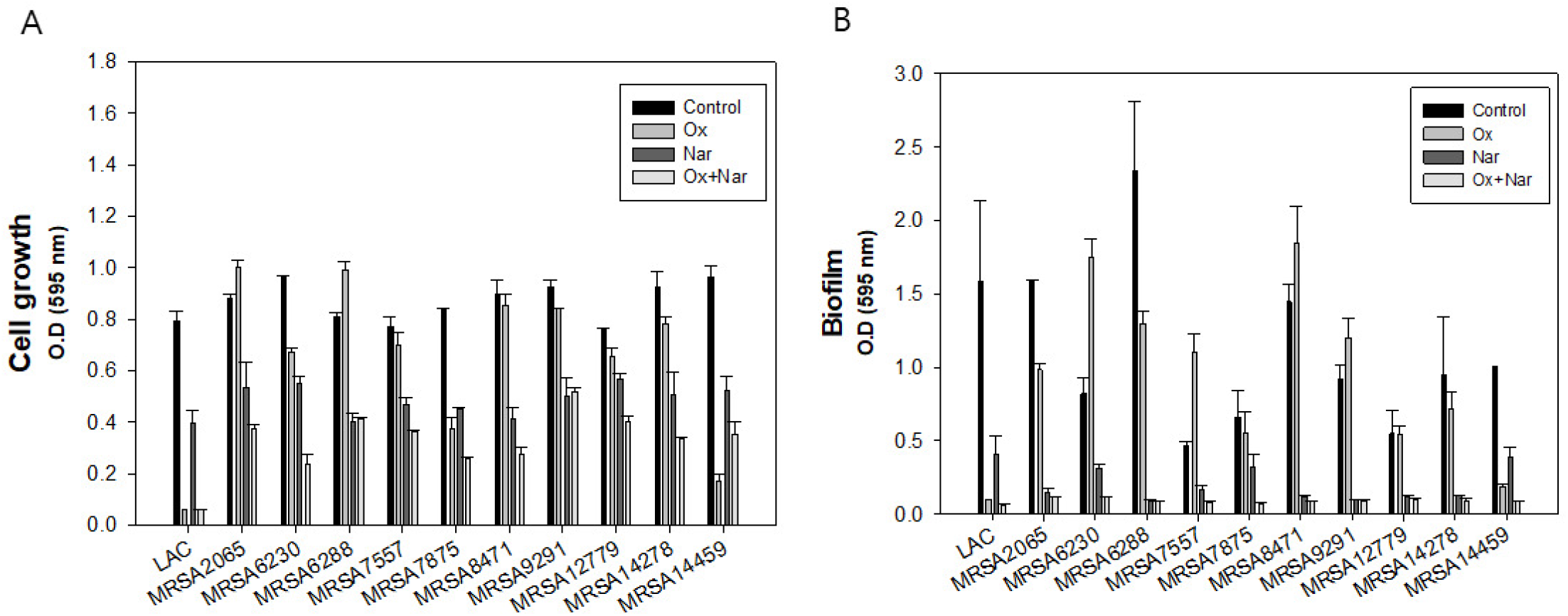
Application of synergetic effect of naringenin and oxacillin to clinically isolated strains. A: Cell growth of LAC and clinical strains with oxacillin and naringenin B: Biofilm formation of LAC and clinical strains with oxacillin and naringenin. The error bars represent the standard deviation of three replicates.

## Discussion

MRSA is notorious for its widespread existence in surgical area and resistance toward antibiotic treatment (20). The overuse of antibiotics for long term had led to the findings of new anti-bacterial agent such as plant derived flavonoids (21). To be more specific, flavonoids are safe as they originated from natural origin and they can be modified to have more efficacy (11). Flavonoids are secondary metabolites which are naturally produced by plant or fungus and they are acting generally as pigment, quorum sensing molecules, antibiotics to other competitive microorganisms (22). Flavonoids have drawn attention from their natural origin and multiple activities which is making bacteria hard to become resistant to flavonoids. For example, they can block respiratory chain, fatty acid synthesis, gyrase activity, quorum sensing molecules and increase membrane permeability etc. (11). However, its functional mechanism on antibacterial activities are not well known. The use of naringenin affected not only distribution of fatty acid in the cell or supernatant but also reducing the expression of biofilm formation genes causing change in biofilm formation and antibiotic sensitivity. Also, cell population became more dispersed when combination therapy was used. As β-lactam antibiotics widely were used, there came out many MRSA strains with much higher or multi-drug resistance. Therefore, there were not many ways of control it but dose dependent treatment of antibiotics. In that point, combinatorial use of flavonoids with antibiotics could be one way to overcome those difficulties especially with higher resistance strains.

## Materials and Methods

### Bacterial strains, media, and culture conditions

For cell preparation, the wild-type strain *Staphylococcus aureus* USA300 (LAC, ATCC^®^ BAA1756) (23) and the mutant strain Δ*agr* (24) were cultured in tryptic soybean broth (TSB) agar and/or liquid broth. For pre-culture, a single colony of the strain from a TSB agar plate was used to inoculate 5 mL of TSB medium. Then, 1% (v/v) of the cell culture suspension was inoculated in a 96-well plate for the antibiotic resistance test and the cell cultivation was conducted overnight in incubator at 37 °C without shaking unless stated otherwise.

### Antibiotic and phytochemical

Oxacillin and a total of 5 selected phytochemical powders from flavonoid group i.e., naringenin, chrysin, daidzein, genistein, apigenin were commercially purchased from Sigma-Aldrich (St. Louis, Minneapolis, USA). Stock solutions of these agents were prepared in sterile dimethyl-sulphoxide (DMSO) solvent to various concentrations.

### Analysis of cell growth and biofilm formation

The cell growth was measured in terms of cell density using a 96-well microplate reader (TECAN, Switzerland). Biofilm formation was analyzed crystal violet staining using the protocols (25). Briefly, supernatant was discarded, and biofilm was fixed with methanol and air dried. Then, biofilm was stained with 200μl of 0.2 % crystal violet solution for 5 min. Next step was to discard crystal violet and wash with distilled water and air-dry step. Finally, biofilm was analyzed at 595 nm using 96-well microplate reader.

### Reverse transcription for cDNA synthesis and semi-quantitative real time PCR

Pre-culture was conducted using 5 ml of TSB with a single colony from agar plate in shaking incubator at 37 °C and 200 rpm during overnight. Cell cultivations were carried out using 5ml of TSB with 1 % inoculum in shaking incubator at 37 °C for 24 h to extract total RNA w/o naringenin or with naringenin addition. Cells were harvested using centrifuge at 3500 rpm for 20 min. Then, total RNA was prepared using TRIzol™ Reagent and reverse transcription was performed with Superscript IV Reverse Transcriptase (Invitrogen Co., Carlsbad, CA) to generate the cDNA following the instructions manuals. Primer design was done using Primer express software v3.0.1 from Thermo Fisher Scientific (Waltham, MA, USA), and these primers can generate 150 bp PCR product for the comparison of gene expressions. Before semi-quantitative PCR, cycle number was optimized to set up the saturated gene expression level of *gyrB* (endogenous control) for each template. After optimization, 25 cycle came out to be the optimal cycle number and further comparative analysis of gene expression became possible. Then, Semi-quantitative PCR was conducted using LA taq with GC buffer I (Takara medical co. ltd) using the methods in manual.

### Fatty acid analysis

Gas chromatography-mass spectrometry (GC-MS) was used for the detection and quantification of total fatty acids and supernatant fatty acids, according to a previously described method with slight modification (26). For total fatty acids analysis, cell cultivation was conducted using 5ml of TSB with 1 % inoculum in shaking incubator at 37 °C and 200 rpm. Then, cell was collected at 24 h and 48 h followed by centrifugation at 3500 rpm for 20 min and washed twice with Milli Q water. Harvested cell was freeze-dried for further methanolysis to analyze total fatty acids. For the analysis of supernatant fatty acids in the medium, spent culture supernatant was incubated in shaking incubator for 2 h at 37 °C and 200 rpm after adding 5 ml of methanol and chloroform to extract fatty acids. The chloroform phase was collected and slowly evaporated under compressed N_2_ at 50 °C in heating block. Fatty acids were re-solubilized with 1 ml of chloroform for further steps. For methanolysis of fatty acids, approximately 10 mg of freeze-dried cells were weighed and placed in Teflon-stoppered glass vials, and then 1 mL chloroform and 1 mL methanol/H_2_SO_4_ (85:15 v/v %) were added to the vials. After incubation at 100°C for 2 h, the vials were cooled to room temperature, and then incubated on ice for 10 min. After adding 1 mL of ice-cold water, the samples were thoroughly mixed by vortexing for 1 min and then centrifuged at 3500 rpm. The organic phases (bottom of the vials) were extracted by a pipette and transferred to clean borosilicate glass tubes containing Na_2_SO_4_. GC-MS was then performed with a Perkin Elmer Clarus 500 gas chromatograph that was connected to a Clarus 5Q8S mass spectrometer at 70 eV (m/z 50-550; source at 230 °C and quadruple at 150 °C) in EI mode with an Elite 5 MS capillary column (30 m × 0.32 mm × 0.25 μm film thickness; J&W Scientific, USA). Helium was used as the carrier gas at a flow rate of 1.0 mL/min. The inlet temperature was maintained at 300 °C, and the oven was programmed to start at 150 °C for 2 min before increasing to 300 °C at a rate of 4 °C/min, and the temperature was maintained for 20 min. The injection volume was 1 μL, with a split ratio of 50:1.

The structural assignments were based on interpretation of the mass spectrometric fragmentation and confirmed by comparison with the retention times and fragmentation patterns of the authentic compounds along with spectral data obtained from the online libraries of Wiley (http://www.palisade.com) and NIST (http://www.nist.gov). The internal standard was 1 μL of methyl heneicosanoate (10 mg/mL) and Bacterial acid methyl ester (BAME) mix (Merck-Millipore, Burlington, MA, USA) was used to identify the each peak of fatty acids and analytical standards for each fatty acid were used for quantification.

## Acknowledgements

This paper was supported by Konkuk University in 2019.

## References

1. Garoy EY, Gebreab YB, Achila OO, Tekeste DG, Kesete R, Ghirmay R, Kiflay R, Tesfu T. 2019. Methicillin-Resistant Staphylococcus aureus (MRSA): Prevalence and Antimicrobial Sensitivity Pattern among Patients-A Multicenter Study in Asmara, Eritrea. Can J Infect Dis Med Microbiol 2019:8321834.

2. Gordon RJ, Lowy FD. 2008. Pathogenesis of methicillin-resistant Staphylococcus aureus infection. Clin Infect Dis 46 Suppl 5:S350–9.

3. Ventola CL. 2015. The antibiotic resistance crisis: part 1: causes and threats. P T 40:277–83.

4. de Tejada GM, Sanchez-Gomez S, Razquin-Olazaran I, Kowalski I, Kaconis Y, Heinbockel L, Andra J, Schurholz T, Hornef M, Dupont A, Garidel P, Lohner K, Gutsmann T, David SA, Brandenburg K. 2012. Bacterial Cell Wall Compounds as Promising Targets of Antimicrobial Agents I. Antimicrobial Peptides and Lipopolyamines. Current Drug Targets 13:1121–1130.

5. Periasamy S, Joo HS, Duong AC, Bach THL, Tan VY, Chatterjee SS, Cheung GYC, Otto M. 2012. How Staphylococcus aureus biofilms develop their characteristic structure. Proceedings of the National Academy of Sciences of the United States of America 109:1281–1286.

6. Lambert PA. 2005. Bacterial resistance to antibiotics: Modified target sites. Advanced Drug Delivery Reviews 57:1471–1485.

7. Chen FJ, Wang CH, Chen CY, Hsu YC, Wang KT. 2014. Role of the mecA Gene in Oxacillin Resistance in a Staphylococcus aureus Clinical Strain with a pvl-Positive ST59 Genetic Background. Antimicrobial Agents and Chemotherapy 58:1047–1054.

8. Panche AN, Diwan AD, Chandra SR. 2016. Flavonoids: an overview. Journal of Nutritional Science 5.

9. Winkel-Shirley B. 2001. Flavonoid biosynthesis. A colorful model for genetics, biochemistry, cell biology, and biotechnology. Plant Physiol 126:485–93.

10. Osonga FJ, Akgul A, Miller RM, Eshun GB, Yazgan I, Sadik OA. 2019. Antimicrobial Activity of a New Class of Phosphorylated and Modified Flavonoids. ACS Omega 4:12865–12871.

11. Gorniak I, Bartoszewski R, Kroliczewski J. 2019. Comprehensive review of antimicrobial activities of plant flavonoids. Phytochemistry Reviews 18:241–272.

12. Lee KA, Moon SH, Lee JY, Kim KT, Park YS, Paik HD. 2013. Antibacterial Activity of a Novel Flavonoid, 7-O-Butyl Naringenin, against Methicillin-Resistant Staphylococcus aureus (MRSA). Food Science and Biotechnology 22:1725–1728.

13. Tsuchiya H. 2015. Membrane Interactions of Phytochemicals as Their Molecular Mechanism Applicable to the Discovery of Drug Leads from Plants. Molecules 20:18923–18966.

14. Vikram A, Jayaprakasha GK, Jesudhasan PR, Pillai SD, Patil BS. 2010. Suppression of bacterial cell-cell signalling, biofilm formation and type III secretion system by citrus flavonoids. J Appl Microbiol 109:515–27.

15. Budzyńska A, Rózalski M, Karolczak W, Wieckowska-Szakiel M, Sadowska B, Rozalska B. 2011. Synthetic 3-Arylidenefl avanones as Inhibitors of the Initial Stages of Biofilm Formation by Staphylococcus aureus and Enterococcus faecalis. Zeitschrift für Naturforschung C, Journal of biosciences 66:104–14.

16. Abreu AC, Serra SC, Borges A, Saavedra MJ, McBain AJ, Salgado AJ, Simoes M. 2015. Combinatorial Activity of Flavonoids with Antibiotics Against Drug-Resistant Staphylococcus aureus. Microb Drug Resist 21:600–9.

17. Amin MU, Khurram M, Khattak B, Khan J. 2015. Antibiotic additive and synergistic action of rutin, morin and quercetin against methicillin resistant Staphylococcus aureus. BMC Complement Altern Med 15:59.

18. Slobodnikova L, Fialova S, Rendekova K, Kovac J, Mucaji P. 2016. Antibiofilm Activity of Plant Polyphenols. Molecules 21.

19. Song H-S, Choi T-R, Han Y-H, Park Y-L, Park JY, Yang S-Y, Bhatia SK, Gurav R, Kim Y-G, Kim J-S, Joo H-S, Yang Y-H. 2020. Increased resistance of a methicillin-resistant Staphylococcus aureus Δagr mutant with modified control in fatty acid metabolism. AMB Express.

20. Chambers HF, Deleo FR. 2009. Waves of resistance: Staphylococcus aureus in the antibiotic era. Nat Rev Microbiol 7:629–41.

21. Xu Z, Li H, Qin X, Wang T, Hao J, Zhao J, Wang J, Wang R, Wang D, Wei S, Cai H, Zhao Y. 2019. Antibacterial evaluation of plants extracts against ampicillin-resistant Escherichia coli (E. coli) by microcalorimetry and principal component analysis. AMB Express 9:101.

22. Koh CL, Sam CK, Yin WF, Tan LY, Krishnan T, Chong YM, Chan KG. 2013. Plant-Derived Natural Products as Sources of Anti-Quorum Sensing Compounds. Sensors 13:6217–6228.

23. CDC. 2003. Outbreaks of community-associated methicillin-resistant Staphylococcus aureus skin infections––Los Angeles County, California, 2002-2003. MMWR Morb Mortal Wkly Rep 52:88.

24. Cheung GYC, Wang R, Khan BA, Sturdevant DE, Otto M. 2011. Role of the accessory gene regulator agr in community-associated methicillin-resistant Staphylococcus aureus pathogenesis. Infection and immunity 79:1927–1935.

25. Shukla S, Toleti SR. 2017. An Improved Crystal Violet Assay for Biofilm Quantification in 96-Well Micro-Titre Plate. doi:10.1101/100214.

26. Bhatia SK, Kim J, Song H-S, Kim HJ, Jeon J-M, Sathiyanarayanan G, Yoon J-J, Park K, Kim Y-G, Yang Y-H. 2017. Microbial biodiesel production from oil palm biomass hydrolysate using marine Rhodococcus sp. YHY01. Bioresource Technology 233:99–109.

